# Alpha-tubulin acetylation in *Trypanosoma cruzi*: a dynamic instability of microtubules is required for replication and cell cycle progression

**DOI:** 10.1101/2020.12.15.422917

**Authors:** Victoria Lucia Alonso, Mara Emilia Carloni, Camila Silva Gonçalves, Gonzalo Martinez Peralta, Maria Eugenia Chesta, Alejandro Pezza, Luis Emilio Tavernelli, Maria Cristina M. Motta, Esteban Serra

## Abstract

Trypanosomatids have a cytoskeleton arrangement that is simpler than what is found in most eukaryotic cells. However, it is precisely organized and constituted by stable microtubules. Such microtubules compose the mitotic spindle during mitosis, the basal body, the flagellar axoneme and the subpellicular microtubules, which are connected to each other and also to the plasma membrane forming a helical arrangement along the central axis of the parasite cell body. Subpellicular, mitotic and axonemal microtubules are extensively acetylated in *Trypanosoma cruzi*. Acetylation on lysine (K) 40 of α-tubulin is conserved from lower eukaryotes to mammals and is associated with microtubule stability. It is also known that K40 acetylation occurs significantly on flagella, centrioles, cilia, basal body and the mitotic spindle in eukaryotes. Several tubulin posttranslational modifications, including acetylation of K40, have been catalogued in trypanosomatids, but the functional importance of these modifications for microtubule dynamics and parasite biology remains largely undefined. The primary tubulin acetyltransferase that delivers this modification was recently identified in several eukaryotes as Mec-17/ATAT, a Gcn5-related N-acetyltransferase. Here, we report that *T. cruzi* ATAT acetylates α-tubulin *in vivo* and is capable of auto-acetylation. *Tc*ATAT is located in the cytoskeleton and flagella of epimastigotes and colocalizes with acetylated α-tubulin in these structures. We have expressed *Tc*ATAT with an HA tag using the inducible vector p*Tc*INDEX-GW in *T. cruzi*. Over-expression of *Tc*ATAT causes increased levels of the acetylated isoform, induces morphological and ultrastructural defects, especially in the mitochondrion, and causes a halt in the cell cycle progression of epimastigotes, which is related to an impairment of the kinetoplast division. Finally, as a result of *Tc*ATAT over-expression we observed that parasites became more resistant to microtubule depolymerizing drugs. These results support the idea that α-tubulin acetylation levels are finely regulated for the normal progression of *T. cruzi* cell cycle.

## 1. Introduction

*Trypanosoma cruzi*, the etiological agent of Chagas disease or American trypanosomiasis, is a kinetoplastid parasite with a complex life cycle that alternates between a mammalian host and an insect host (Triatominidae family), which is the biological vector of this disease. The world Health Organization classifies Chagas disease as one of the 13 most neglected tropical diseases, constituting a very important social and economic problem in Latin America (WHO, 2012; http://who.int). Trypanosomatids have a cytoskeleton arrangement that is simpler than what is found in most eukaryotic cells. However, it is precisely organized and constituted by stable microtubules (MT). Such MTs are present in the mitotic spindle during mitosis, the basal body, the flagellar axoneme and the subpellicular MTs, which are connected to each other and also to the plasma membrane, thus forming a helical arrangement along the central axis of the parasite cell body (Vidal et al., 2017). MTs provide the basis for cytoskeletal architecture and are formed by α/β-tubulin heterodimers, comprising 13 typical protofilaments connected to each other forming helical tubes. These structures are regulated by interacting with a variety of MT-associated proteins (MAPs), also by a differential expression of α and β-tubulin genes (tubulin isotypes) and by a plethora of post-translational modifications (PTMs) (Gadadhar et al., 2017). Several conserved lysines in α- and β-tubulin are acetylated in eukaryotes, and acetylation of the α-tubulin luminal residue lysine 40 (K40) has been the most characterized since its discovery over thirty years ago (L’Hernault and Rosenbaum, 1985; Al-Bassam and Corbett, 2012; Kull and Sloboda, 2014; Eshun-Wilson et al., 2019). Acetylation of α-tubulin on K40 was associated with the stability of microtubules and described as a marker of microtubules resistance to depolymerizing drugs from the beginning of its study (L’Hernault and Rosenbaum, 1985; Piperno et al., 1987). In most cells acetylated α-tubulin is a minor isoform, observed in primary cilia, flagella, centrioles and neuronal axons (Piperno et al., 1987; Hubbert et al., 2002; Kalebic et al., 2012; Nakakura et al., 2015). In contrast, trypanosomatids have a significantly high proportion of acetylated α-tubulin, concentrated in the subpellicular, mitotic and axonemal MTs (Sasse and Gull, 1988; Souto-Padron et al., 1993), which makes these organisms attractive models to study the function of α-tubulin K40 acetylation.

Although acetylation typically correlates with stable and long-lived microtubules in cells, acetylation itself does not confer stability, but may rather make microtubules more resilient to mechanical forces (Howes et al., 2013; Szyk et al., 2014; Coombes et al., 2016; Portran et al., 2017). Yet despite years of study, the effects of acetylation on MTs and MT function in cells are still debated. The primary α-tubulin acetyltransferase that delivers this modification was recently identified in several eukaryotes as MEC-17/ATAT, a Gcn5-related N-acetyltransferase containing a catalytic domain that is conserved from protists to mammalian species. MEC-17/ATAT directly promotes α-tubulin acetylation *in vitro* and it is the major α-tubulin acetyltransferase *in vivo* (Akella et al., 2010; Shida et al., 2010). MEC-17 is required for touch sensation in *Caenorhabditis elegans*, normal embryonic development in zebrafish, and the rapid assembly of primary cilia in RPE-hTERT cells (Akella et al., 2010; Shida et al., 2010; Li et al., 2012). Also, acetylation does not seem to be only a passive mark on microtubules, as its loss disrupts microtubule structural integrity in touch receptor neurons, leading to axonal morphology defects (Cueva et al., 2012). Loss of ATAT also causes brain abnormalities in mice (Kim et al., 2013). ATAT was characterized in the apicomplexan parasite *Toxoplasma gondii* where it was shown that K40 acetylation stabilizes MTs and is required for replication. *Tg*ATAT is expressed in a cell cycle-regulated manner and genetic disruption ablates K40 acetylation, thus inducing replication defects, since parasites appear to initiate mitosis but exhibit an incomplete or improper nuclear division (Varberg et al., 2015).

Several tubulin PTMs, including acetylation of K40, have been catalogued in trypanosomatids (Rosenzweig et al., 2008; Nett et al., 2009; Tsigankov et al., 2013; Moretti et al., 2018), but the functional importance of these modifications for MT dynamics and parasite biology remains largely undefined. We have studied the effect of α-tubulin hyperacetylation on *T. cruzi* cell cycle by over-expressing its α-tubulin acetyltransferase (*Tc*ATAT) using the tetracycline-inducible vector p*Tc*INDEX-GW (Alonso et al., 2014a). This system allowed us to control the amount of *Tc*ATAT, and hence the amount of acetylated α-tubulin in epimastigotes. Over-expressing parasites showed an increase of acetylated α-tubulin as expected that was associates to growth defects related to a cell cycle arrest and impairment of kinetoplast division. *Tc*ATAT is located in the cytoskeleton and flagella of *T. cruzi* and colocalizes with acetylated α -tubulin. Over-expression also induced morphological alterations, that are related to cell division impairment, and ultrastructural changes, especially in the mitochondrial branches and in kDNA topology. These evidence supports the idea that α-tubulin acetylation is tightly regulated in *T. cruzi* and indicates that although the cytoskeleton arrangement is considered stable in trypanosomatids, a dynamic instability of microtubules is required for replication and cell cycle progression.

## 2. Materials and Methods

### 2.1. Molecular cloning of TcATAT-HA

*Tc*ATAT gene from *T. cruzi* Dm28*c* strain were amplified using the following oligonucleotides, TATFw:AA<u>GGATTC</u>**ATGTATCCGTATGATGTCCCGGATTATGCT**AGTTCCACATCGCAA and TATRv:AA<u>CTCGAG</u>TGTTCTGGAGTACCACT, adding and HA-tag in the N-terminus (in bold). DNA purified from *T. cruzi* Dm28*c* epimastigotes was used as template. The PCR products obtained with a proofreading DNA polymerase were inserted into pCR2.1-TOPO vector (Invitrogen) and sequenced. *Tc*ATAT-HA coding regions was then inserted into a pENTR3C vector (Gateway system Invitrogen) using the *Bam*HI/*Xho*I restriction sites included in the oligonucleotides (underlined) and then transferred to pDEST17 (Gateway system Invitrogen) and p*Tc*INDEX-GW vectors by recombination using LR clonase II enzyme mix (Invitrogen). The pDEST17 constructs were transformed into *Escherichia coli* BL21 pLysS and recombinant proteins, fused to a six histidine-tag, were obtained by expression-induction with 0.5 mM IPTG for 3 h at 30°C. The proteins were purified by affinity chromatography using a Ni-NTA agarose resin (Qiagen) following the manufacturer’s instructions.

### 2.2. *Trypanosoma cruzi* culture and transfection

*T. cruzi* Dm28*c* epimastigotes were cultured at 28 °C in LIT medium (5 g/L liver infusion, 5 g/L bacto-tryptose, 68 mM NaCl, 5.3 mM KCl, 22 mM Na2HPO4, 0.2% (w/v) glucose and 0.002% (w/v) hemin) supplemented with 10% (v/v) heat-inactivated, UV-irradiated Fetal Calf Serum (FCS) (Internegocios S.A, Argentina). Viability was determined by counting live cells with a haematocytometer using Erythrosin B staining. For half media inhibitory concentration (IC_50_) calculations parasites where treated with Oryzalin (0-300 μM) for 72 h. and the number of parasites was plotted against the log[Oryzalin]. The plot was fitted with the non-parametric regression log(inhibitor) vs. response -Variable slope (four parameters) in GraphPad Prism version 8.0. Epimastigotes’ motility was examined using the computer-assisted semen analysis (CASA) system (Microptic, SCA evolution). Parameters used were as follows: 30 frames acquired, frame rate of 60 Hz, and cell size of 10–100 μm^2^. At least 30 microscopy fields corresponding to a minimum of 300 epimastigotes were analyzed in each experiment.

Epimastigotes from *T. cruzi* Dm28*c* were transfected with the pLEW13 plasmid to generate parasites expressing T7 RNA polymerase and the tetracycline repressor using a nucleofection method. Briefly, epimastigotes were cultured in LIT medium at 28 °C to a final concentration of 4×10^7^ parasites per transfection. Then, parasites were harvested by centrifugation at 1500 g for 10 min at room temperature, washed once with phosphate buffered saline (PBS) and resuspended in 0.4 mL BSF transfection buffer (5 mM KCl, 0.15 mM CaCl_2_ 90 mM, Na_2_HPO_4_ 50 mM HEPES pH 7.3). Nucleofection (Nucleofector 2B, Lonza) was performed in a 0.2 cm gap cuvette (Bio-Rad) with ∼20 μg of plasmid DNA added to a final volume of 400 μL. The parasite-DNA mixture was kept on ice for 20 min prior to nucleofection with program X-014. After nucleofection, cells were transferred into 3 mL of LIT medium containing 20% FCS, maintained at room temperature for 15 minutes and then incubated at 28 °C. Geneticin (G418; Life Technologies) was added at a concentration of 200 μg/mL, and parasites were incubated at 28 °C. After selection, pLEW13 transfected epimastigotes were maintained in the presence of 100 μg/ml of G418 (Sigma Aldrich). This parental cell line was then nucleofected with p*Tc*INDEX-GW *Tc*ATAT-HA construct following a similar protocol and transgenic parasites were obtained after 4 weeks of selection with 100 μg/ml G418 and 200 μg/ml Hygromycin B (Sigma Aldrich).

### 2.3. Polyclonal antibodies

All experiments were approved by the Institutional Animal Care and Use Committee of the School of Biochemical and Pharmaceutical Sciences, National University of Rosario (Argentina) (File 6060/227) and conducted according to specifications of the US National Institutes of Health guidelines for the care and use of laboratory animals. Rabbits were only used for the production of polyclonal antibodies. A rabbit was immunized two times with recombinant ATAT-HA protein purified from *E. coli* and an equal volume of Freund’s adjuvant. The animal was bled two weeks after the final injection.

### 2.4. Protein extracts

Exponentially growing epimastigotes were washed twice with cold PBS, and the pellets were resuspended in lysis buffer (20 mM HEPES, 8 M Urea) and incubated for 30 min at room temperature with gentle agitation. Insoluble debris was eliminated by centrifugation. The same procedure was applied to amastigote and trypomastigote cellular pellets. *T. cruzi* cytoskeleton-enriched extracts were prepared as previously described (Alonso et al., 2014b).

### 2.5. Western blot

Protein extracts were fractioned in SDS-PAGE and transferred to a nitrocellulose membrane. Transferred proteins were visualized with Ponceau S staining. Membranes were treated with 10% non-fat milk in PBS for 2 hours and then incubated with specific antibodies diluted in 0.5% Tween 20 in PBS (PBS-T) for 3 hours. Primary antibodies used were: rat monoclonal anti-HA 1:2000 (ROCHE), affinity-purified rabbit polyclonal anti-*Tc*ATAT 1:200, mouse monoclonal anti-trypanosome α-tubulin clone TAT-1 1:1000 (a gift from K. Gull, University of Oxford, UK), rabbit polyclonal anti-Acetyl-lysine 1:1000 (Millipore), mouse monoclonal anti-acetylated α-tubulin clone 6-11B-1 1:2000 (Sigma Aldrich). Bound antibodies were detected using peroxidase-labeled anti-rabbit IgG (GE Healthcare), anti-mouse IgG (GE Healthcare) or anti-rat IgG (Thermo Scientific) and developed using ECL Plus kit (GE Healthcare) according to manufacturer’s protocols. Immunoreactive bands were visualized and photographed in the Amersham Imager 600 (GE Healthcare). Images were processed and bands where quantified with ImageJ (Miller, 2010).

### 2.6. Preparation of cytoskeletal and flagellar complexes

The isolated cytoskeletons and flagellar complexes were obtained as previously described (Alonso et al., 2016) and followed by the immunofluorescence protocol as described below.

### 2.7. Immunofluorescence

Trypomastigotes and exponentially growing epimastigotes were centrifuged, washed twice in PBS, settled on polylisine-coated (Sigma Aldrich) coverslips and fixed with 4% para-formaldehyde in PBS at room temperature for 20 minutes. For the mitochondrial staining, epimastigotes were resuspended in PBS and incubated with 1 μM MitoTracker Orange CMTMRos (Invitrogen) for 30 minutes at 28°C, washed twice in PBS and fixed with 4% para-formaldehyde. Fixed parasites were washed with PBS and permeabilized with 0.1% Triton X-100 in PBS for 10 minutes. After washing with PBS, parasites were incubated with the appropriate primary antibody diluted in 5% BSA in PBS for 2 hours at room temperature. Primary antibodies used were: rat monoclonal anti-HA 1:200 (ROCHE), affinity-purified rabbit polyclonal anti-*Tc*ATAT 1:20, mouse monoclonal anti-acetylated α-tubulin clone 6-11B-1 1:100 (Sigma Aldrich) and mouse polyclonal anti-PAR2 1:100 (*T. cruzi* paraflagellar rod 2 protein). In colocalization experiments both antibodies were incubated together. Non-bound antibodies were washed with 0.01% Tween 20 in PBS and then the slides were incubated with fluorescent-conjugated anti-mouse Alexa-555 (Invitrogen) or anti-rat (FITC, Invitrogen) and anti-rabbit (FITC, Jackson Immuno Research) IgG antibodies and 2 μg/mL of DAPI for 1 hour. The slides were washed with 0.01% Tween 20 in PBS and finally mounted with VectaShield (Vector Laboratories). Images were acquired with a confocal Zeiss LSM880 and Nikon Eclipse Ni-U epifluorescence microscope. ImageJ software were used to process all images.

### 2.8. Ultrastructural analysis

#### 2.8.1. Scanning electron microscopy (SEM)

Cells were fixed in 2.5% glutaraldehyde diluted in 0.1 M cacodylate buffer (pH 7.2) for 1 h, following washes in the same buffer and then adhered to poly-L-lysine-coated microscope coverslips. After fixation, parasites were post-fixed with 1% osmium tetroxide diluted in cacodylate buffer for 1 hour, dehydrated in ethanol (50%, 70%, 90%, and two exchanges of 100%, 10 min in each step), critical point dried in CO_2_ by using a Leica EM CPD030 equipament (Leica, Wetzlar, Germany) and ion sputtered in a Balzers FL9496 unit (Postfach 1000 FL-9496 Balzers Liechtenstein). Samples were observed under an EVO 40 VP SEM (Zeiss, Germany).

#### 2.8.2. Transmission electron microscopy (TEM)

Cells were fixed as described for Scanning electron microscopy. Then, samples were post-fixed in 1% osmium tetroxide and 0.8% potassium ferricyanide, diluted in the same buffer, for 1 h. After this, parasites were washed in cacodylate buffer, dehydrated in a graded series of acetone (50%, 70%, 90%, and two exchanges of 100%, 10 min in each step) and embedded in Polybed resin (Epon) (Electron Microscopy Sciences, Hatfield, PA, USA). Ultra-thin sections were stained with uranyl acetate for 40 min and then with lead citrate for 5 min. Samples were observed under a Jeol 1200 EX TEM operating at 80 kV (Jeol, Japan).

### 2.9. Cell Cycle analysis

Synchronization of epimastigotes in G1 of the cell cycle was achieved using hydroxyurea (HU). Cells in exponential growth phase were arrested by incubation with 20 mM of HU for 24 h and then released by washing twice with PBS and suspending the cells in culture medium. Cells continued to be cultured for 24 h and samples were taken at the indicated time points. Cell cycle progression of parasites was analyzed by flow cytometry as described previously (Tavernelli et al., 2019). Briefly, one million cells were fixed with cold 70% ethanol and then washed with PBS and stained with 20 μg/ml Propidium Iodide (PI) in buffer K (0.1% sodium citrate, 0.02 mg/mL RNAse A (Sigma), and 0.3% NP-40). Ten thousand events per sample were acquired using BD Cell Sorter BD FACSAria II. Results were analyzed with FlowJo software.

### 2.10. ATAT-HA purification form *T. cruzi* epimastigotes

A culture of 50 ml of *T. cruzi* Dm28*c* p*Tc*INDEX-GW-ATAT-HA induced with 0.5 μg/mL tetracycline for 24 hours was collected and resuspended in 500 μL lysis buffer MME (Mops pH 6,9 10 mM, EGTA-EDTA 1 mM, MgSO_4_ 1 mM) supplemented with NaCl 1M, Triton X-100 0.2% and protease inhibitor cocktail (GE). Incubated with agitation at 4°C for 30 min and then cells were ruptured by sonication and centrifuged for 20 min. at 16,000 g. The supernatant was loaded to an anti-HA agarose column (Roche) following the manufacturers’ instructions. The flowthrough was collected, and the column was washed with 1 volume of PBS 0.5% Tween-20. The bound protein was eluted four times with 250 μL of HA peptide (Sigma-Aldrich) (0,1 mg/ml).

### 2.11. Autoacetylation assay

0.5 μg of ATAT-HA purified from *T. cruzi* epimastigotes was incubated in the absence or presence of 0.5 mM Acetyl CoA (Sigma) for 1.5 h at 37°C in acetylation buffer (50mM Tris–HCl, pH 8.0; 10% glycerol; 1mM MgCl_2_; 1mM DTT; 1mM PMSF; 20mM sodium butyrate). The samples were resolved on SDS-PAGE gels, transferred to nitrocellulose membranes (GE Healthcare) and subjected to western blot analysis as described above.

## 3. Results

### 3.1. ATAT homologue in *T. cruzi* is expressed in all *T. cruzi* life cycle stages

A bioinformatic survey of the *T. cruzi* genome in TriTrypDB (https://tritrypdb.org/) revealed a single gene containing a MEC-17 domain belonging to the Gcn5-related superfamily (PF05301), described in this database as a putative alpha-tubulin N-acetyltransferase. In the genome of the Dm28*c* strain the predicted protein sequence is 330 amino-acids long with the acetyltransferase domain in its N-terminal portion (C4B63_12g332), from now on we will name it *Tc*ATAT (Figure 1). When we looked for homologs in other trypanosomatids, we found that the in *T. brucei* (Tb927.3.1400) the putative alpha-tubulin N-acetyltransferase contains 55% of identical residues compared to *T. cruzi* and that this homology is widespread along the sequence. In the case of *Leishmania mayor* (LmjF.25.1150) the identity is restricted to the acetyltransferase domain (50% identical residues) (Supplementary Figure S1A). When we aligned the acetyltransferase domains of ATAT homologues from several representative species with Clustal Omega, we observed that the acetyltransferase domain is highly conserved, including the key residues critical for the enzymatic activity (asterisks in Figure 1B), while both N-terminal and C-terminal portions of the different homologs are highly dissimilar and even have different lengths (Figure 1A). For example, the *Toxoplasma gondii* ATAT is considerably larger than all previously characterized ATAT/MEC-17 proteins (Varberg et al., 2015).

**Figure 1:**
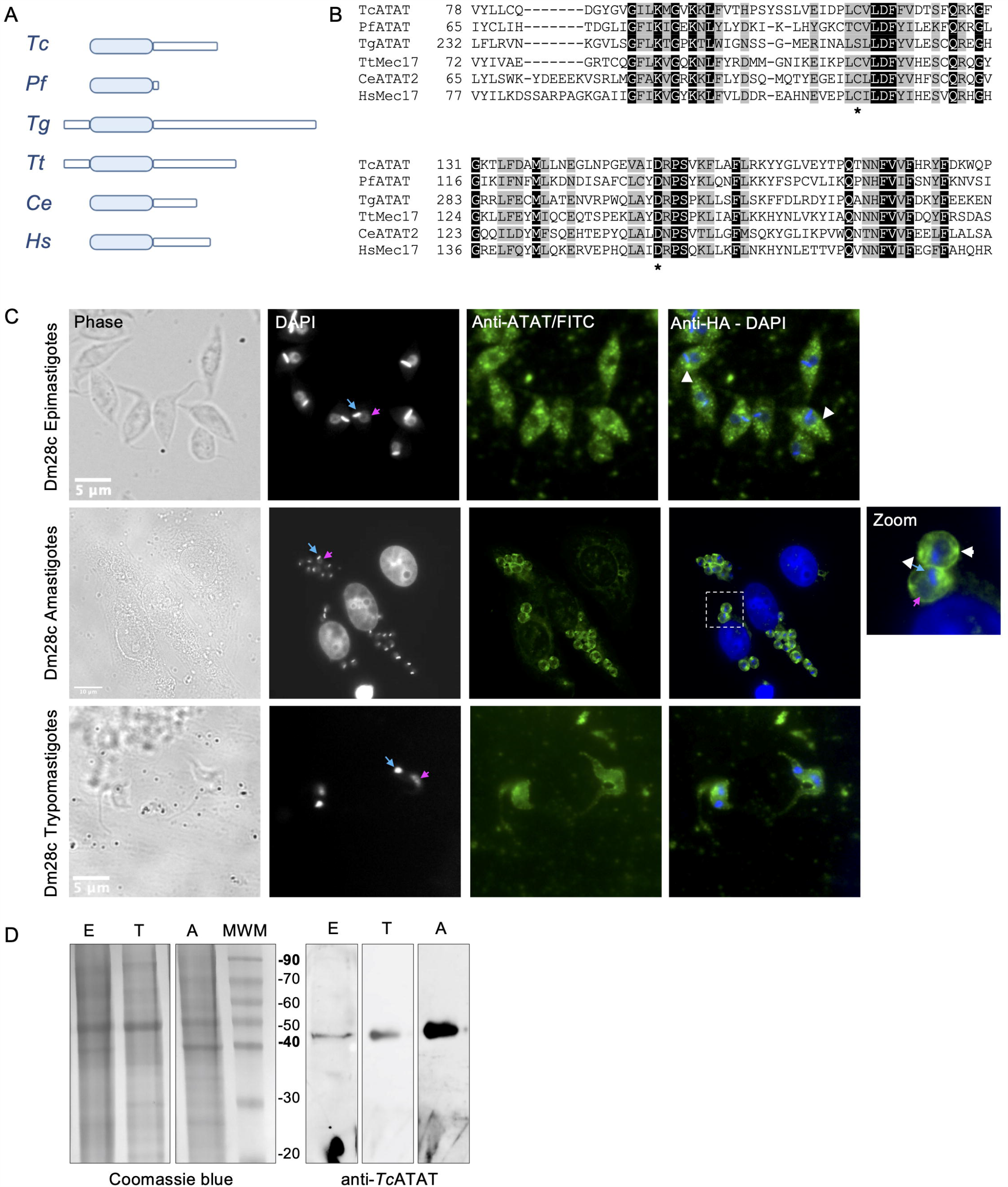
The GCN5 acetyltransferase domain is conserved in *T. cruzi*. **(A)** Schematic representation of ATAT/Mec17 form different organisms (in order: *T. cruzi, Plasmodium falciparum, Toxoplasma gondii, Tetrahymena thermophila, Caenorhabditis elegans, Homo sapiens*). The acetyltransferase domain is represented as light blue rectangles. **(B)** Multiple sequence alignment of the acetyltransferase domain using T-coffee and colored with Boxshade. Two conserved residues important for catalysis in *Hs*ATAT are marked with asterisks. ***Tc*ATAT is expressed in al life cycle stages of *T. cruzi*. (C)** Immunolocalization of *Tc*ATAT in Dm28c epimastigotes, amastigotes and trypomastigotes using rabbit polyclonal anti-*Tc*ATAT antibodies. Bar: 10 μm. DAPI was used as nucleus and kinetoplast marker. The light blue arrow indicates the kinetoplast and the pink arrow indicates the nucleus. **(D)** Total extracts of Dm28c epimastigotes (E), trypomastigotes (T) and amastigotes (A) were separated by SDS/PAGE and stained with Coomassie Blue (left panel), followed by western blot analysis using rabbit monoclonal anti-*Tc*ATAT antibodies (right panel).

RNAseq experiments showed that *Tc*ATAT transcripts are more abundant in trypomastigote stage compared to epimastigote stage (Smircich et al. 2015) and that they peak in late amastigote and trypomastigote stages (Li et al., 2016). To assess the expression pattern of *Tc*ATAT in all the three stages of *T. cruzi* life cycle we obtained polyclonal antibodies from rabbit against the recombinant protein that were purified and used in western blot and immunofluorescence assays in epimastigotes, amastigotes and trypomastigotes. We observed that *Tc*ATAT is expressed in all life cycle stages, located in the whole cell body of *T. cruzi* and apparently excluded from the nuclei (Figure 1C). Some discrete spots in the perinuclear area in epimastigotes and amastigotes were also observed (Figure 1C, white arrowheads). As expected, we detected *Tc*ATAT by western blot in whole extracts of epimastigotes, trypomastigotes and amastigotes, with a higher expression in the last stage (Figure 1D).

### 3.2. *Tc*ATAT-HA has acetyltransferase activity and acetylates α-tubulin in the cytoskeleton and flagellum of epimastigotes

To determine the impact of *Tc*ATAT on α-tubulin acetylation we obtained *T. cruzi* epimastigotes stably transfected with the p*Tc*INDEX-GW vector (Alonso et al., 2014a) baring the ATAT coding sequence with a hemagglutinin tag on its N-terminus (HA). This plasmid allowed us to induce the expression of the transgene with tetracycline (Taylor and Kelly, 2006). We corroborated the over-expression of *Tc*ATAT-HA in epimastigotes by immunofluorescence and western blot assays with anti-HA antibodies 24 h post-induction (p.i.) (Figure 2A and 2B), and we did not detect tagged protein without tetracycline induction (Figure 2B and Supplementary Figure S2). At this time, an evident phenotypic defect was detected in the induced parasites (Figure 2A, arrowhead) and a round refringent structure was observed in a proportion of the over-expressing epimastigotes.

Then, we quantified de amount of acetylated α-tubulin at different induction times by densitometry (Figure 2C) in the over-expressing epimastigotes. The amount of acetylated α-tubulin increased with *Tc*ATAT-HA induction time being almost 10 times higher at 24 h.p.i. compared to the uninduced control. ATAT shows autoacetylation activity in other organisms (Kalebic et al., 2012; Zhou et al., 2018) and *Tc*ATAT was predicted to be acetylated on K263 with the PAIL server (Deng et al., 2016). To study if this was also the case for *T. cruzi* we performed an autoacetylation assay using purified *Tc*ATAT-HA from epimastigotes incubated in the absence or presence of Acetyl-CoA. We detected acetylated *Tc*ATAT only in the presence of Acetyl-CoA confirming that *Tc*ATAT has a fully functional domain with acetyltransferase activity (Figure 2D).

**Figure 2:**
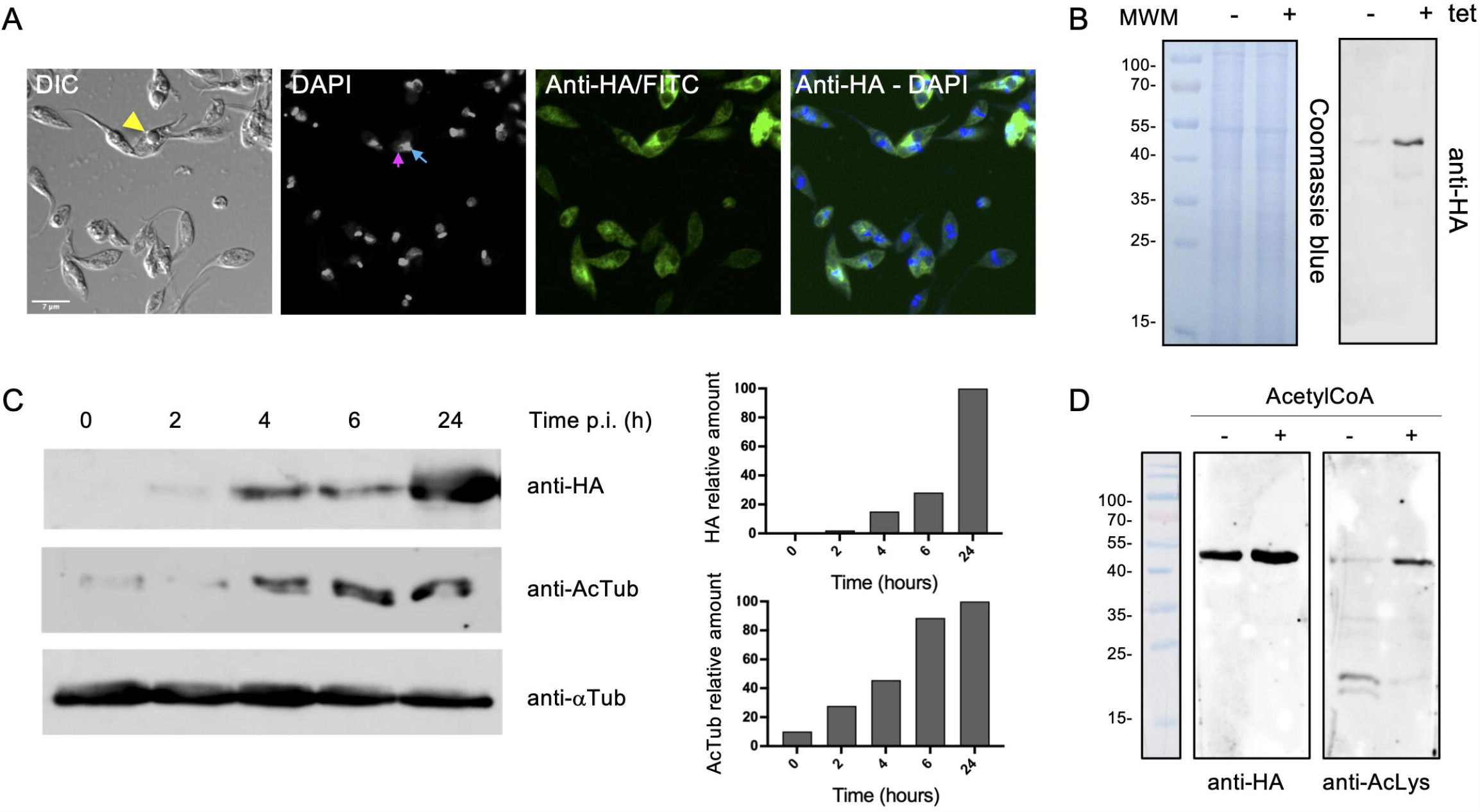
Over-expression of ATAT-HA in epimastigotes increases α-tubulin acetylation. **(A)** Immunolocalization of ATAT-HA with rat monoclonal anti-HA antibodies in Dm28c p*Tc*INDEX-GW ATAT-HA epimastigotes induced with 0.5 μg/ml tetracycline for 24 h. Bar: 5 μm. DAPI was used as nucleus and kinetoplast marker. The light blue arrow indicates the kinetoplast and the pink arrow indicates the nucleus. **(B)** Total extracts of p*Tc*INDEX-GW ATAT-HA epimastigotes in the absence (-) or presence (+) of 0.5 μg/ml tetracycline for 24 h were separated by SDS/PAGE and stained with Coomassie Blue (left panel), followed by western blot analysis using rat monoclonal anti-HA antibodies (right panel). **(C)** Western blot of total extracts of p*Tc*INDEX-GW ATAT-HA epimastigotes with 0.5 μg/ml tetracycline a different time points post-induction using rat monoclonal anti-HA, mouse monoclonal anti-acetylated a-tubulin (anti-AcTub), rabbit monoclonal anti-*Tc*ATAT antibodies and mouse monoclonal anti-a-tubulin (anti-αTub). Bands were quantified by densitometry using α-tubulin signal to normalize the amount of ATAT-HA and acetylated α-tubulin (right panel). **(D)** ATAT-HA autoacetylation assay. ATAT-HA was purified from *T. cruzi* epimastigotes and incubated in the absence (-) and presence (+) of AcetylCoA and then separated by SDS/PAGE followed by western blot analysis with rat monoclonal anti-HA antibodies and rabbit monoclonal anti-Acetylated Lysine (anti-AcLys).

We also tested whether ATAT-HA overexpressing epimastigotes were more resistant to the microtubule-disrupting drug Oryzalin to correlate acetylation with stability. Oryzalin effect on *T. cruzi* has not been reported yet, but we found in the literature that a similar dinitroaniline, Trifuralin, had an IC_50_ between 70 and 160 µM depending on the strain used (Traub-Cseko et al., 2002). To begin with, we determined Oryzalin IC_50_ in Dm28*c* epimastigotes (250.5 μM) and found that it was higher than what was reported for Trifuralin (Supplementary Figure S3A). Epimastigotes treated with Oryzalin show a dose-dependent loss of normal morphology: at higher concentrations epimastigotes adopt a rounded shape with a shorter flagellum - perhaps due to alterations in the polymerization of axonemal microtubules (Supplementary Figure S3B). In presence of 200 μM Oryzalin, Dm28*c* p*Tc*INDEX-GW ATAT-HA epimastigotes induced with tetracycline grew better than uninduced parasites (Table I), as expected for an increased amount of acetylated α-tubulin.

**Table I:** Dm28c pTcINDEX ATAT-HA relative growth in the absence and presence of 200 mM Oryzalin for 72 hours. ** p<0.05 (t-student’s test).

| Dm28c pTcINDEX ATAT-HA relative growth | Tet - | Tet + |
| --- | --- | --- |
| Without drug | 100% | 100% |
| 200 mM Oryzalin | 69% | 85% (**) |

To better characterize the localization of *Tc*ATAT, we isolated subpellicular microtubules and flagellar complexes from transfected epimastigotes and analyzed the presence of *Tc*ATAT-HA in these structures. As observed in Figure 3A, ATAT-HA colocalizes with acetylated α-tubulin in the subpellicular microtubules and the flagellar axoneme. The round structured observed by light microscopy remains insoluble after treatment with detergent and NaCl and is labeled with the anti-HA/FITC antibody, When the confocal images of these preparations are not over-exposed (necessary to observe the cytoskeletal labelling, which is weaker than in the round structure) it appears that ATAT-HA is surrounding this structure (Supplementary Figure S4A). In intact epimastigotes fixed with para-formaldehyde confocal microscopy *Tc*ATAT-HA was observed as accumulated in the periphery of these round structures (Figure 2A). We performed a Z-stack confocal imaging and 3D reconstruction (Supplementary movie V1) and observed that *Tc*ATAT-HA formed a ball-like structure near the nucleus and the kinetoplast. Furthermore, when we compare cytoskeletal preparations of epimastigotes uninduced and induced with tetracycline we observed a disruption of the acetylated MTs upon ATAT-HA over-expression (Supplementary Figure S4B). As a control we verified that *Tc*ATAT-HA did not co-localize with the paraflagellar rod that runs parallel to the axoneme (Supplementary Figure S4C). Also, we obtained protein extracts enriched in cytoskeletal and flagellar proteins and observed by western blot that both the endogenous ATAT and the over-expressed version are only present in the fraction that corresponds to insoluble cytoskeletal and flagellar proteins (P, in figure 3B), confirming that it is tightly associated to these structures.

**Figure 3:**
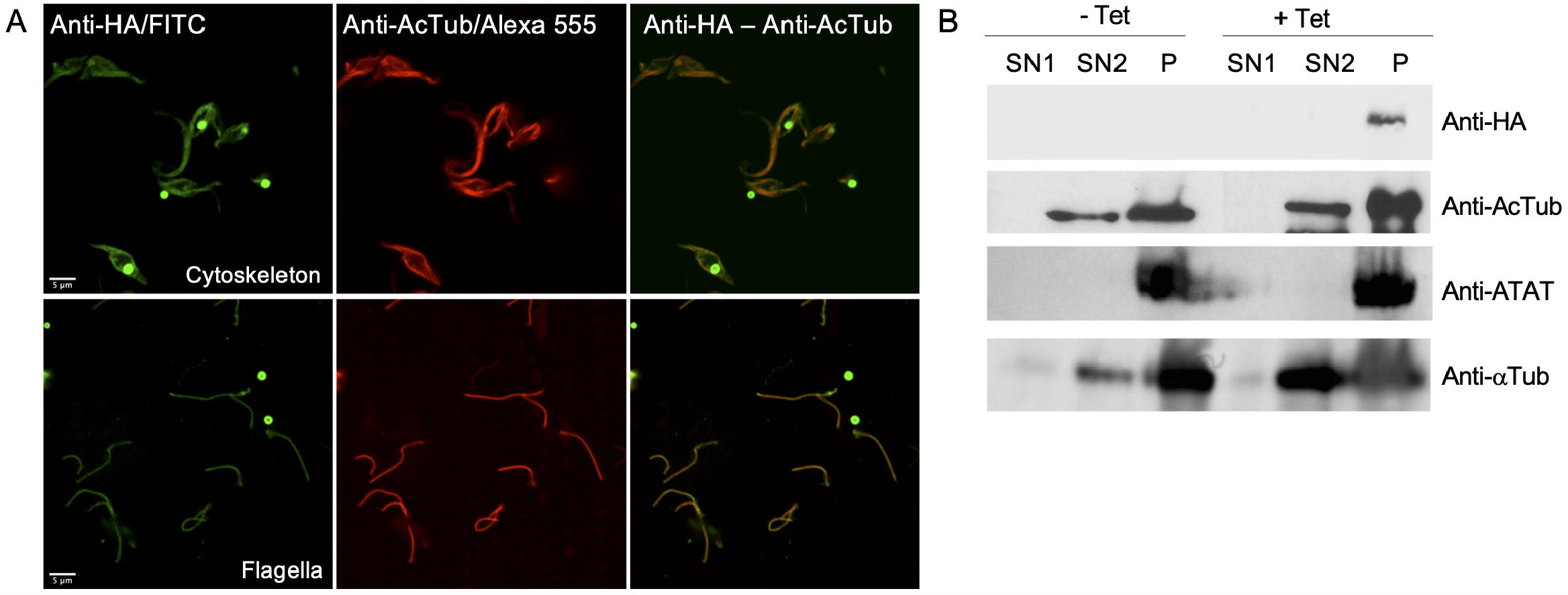
ATAT-HA colocalizes with acetylated α-tubulin in the cytoskeleton and flagella of epimastigotes. **(A)** Immunolocalization of ATAT-HA with rat monoclonal anti-HA antibodies and mouse monoclonal anti-acetylated α-tubulin (anti-AcTub) in isolated cytoskeletons and flagella of Dm28c p*Tc*INDEX-GW ATAT-HA epimastigotes induced with 0.5 μg/ml tetracycline for 24 h. **(B)** Extracts enriched in cytoskeletal and flagellar proteins were analyzed by western blot with rat monoclonal anti-HA antibodies, rabbit polyclonal anti-*Tc*ATAT antibodies, mouse monoclonal anti-acetylated α-tubulin (anti-AcTub) and anti α-tubulin (anti-αTub). SN1, soluble protein extracts; SN2, soluble cytoskeletal and flagellar protein extracts; P, insoluble cytoskeletal and flagellar protein extracts.

The refringent button-like structure observed in the over-expressing parasites was quantified and it was present in approximately 20% of the epimastigotes 48 h.p.i. We also determined that this structure grew with induction time and was usually observed near the nucleus and the kinetoplast (Figure 4A). Transmission Electron Microscopy (TEM) analyses revealed that this round structure is electrodense, not delimited by membrane, and sometimes is seen in continuity with the endoplasmic reticulum, resembling an inclusion body (Figure 4B). These results correlate with the accumulation of *Tc*ATAT-HA observed in association to the isolated cytoskeleton where the structure is seen connected to the flagellum (Figure 3A).

**Figure 4:**
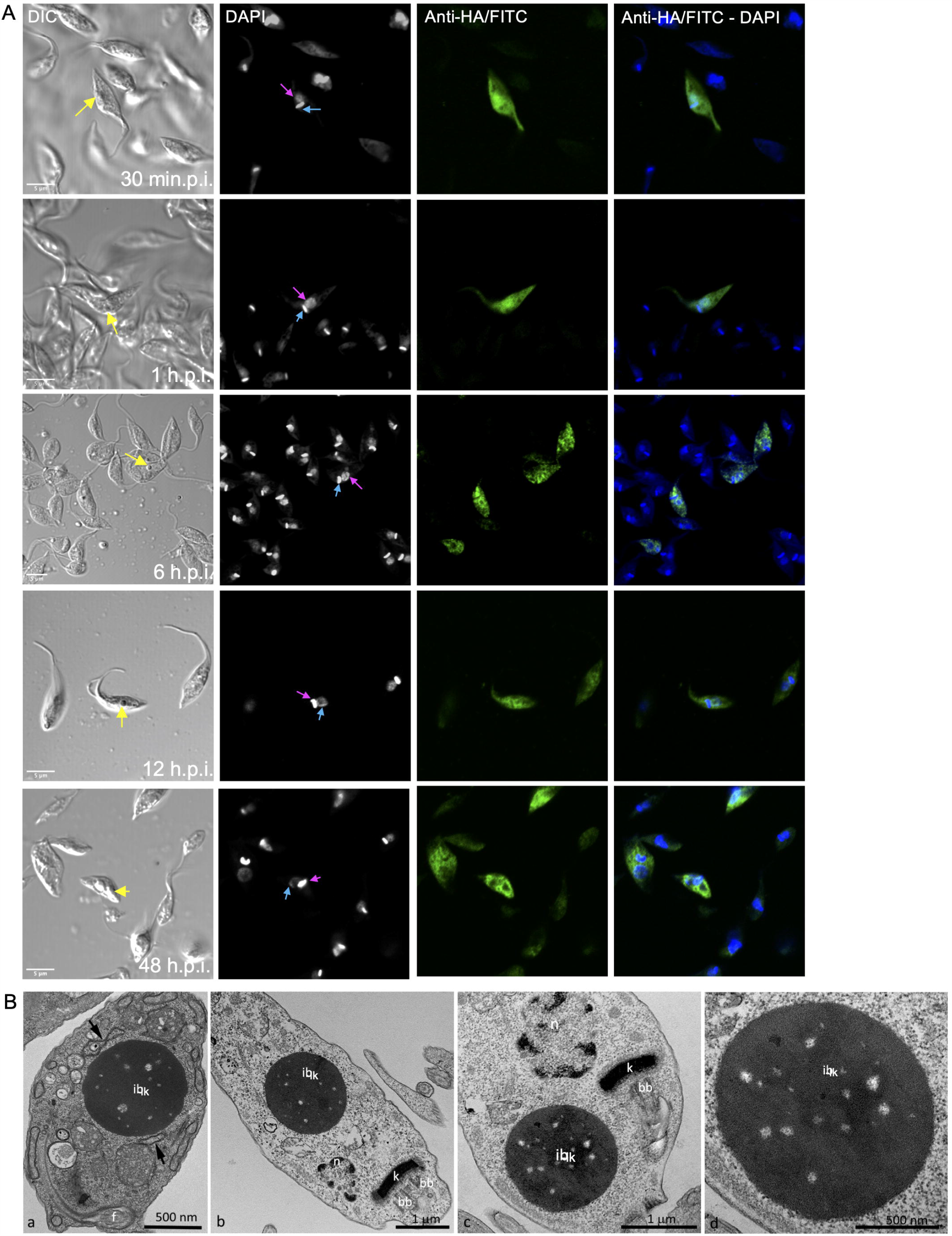
Over-expression of ATAT-HA induced the formation of an inclusion body-like structure. **(A)** Immunolocalization of ATAT-HA with rat monoclonal anti-HA antibodies in Dm28c p*Tc*INDEX-GW ATAT-HA epimastigotes induced with 0.5 mg/ml tetracycline at different time points p.i. The yellow arrow indicates the inclusion body-like structure, the light blue arrow indicates the kinetoplast and the pink arrow indicates the nucleus. DAPI was used as nucleus and kinetoplast marker. Bar: 5 μm. **(B)** Transmission Electron Microscopy of Dm28c p*Tc*INDEX-GW ATAT-HA epimastigotes induced with 0.5 μg/ml tetracycline for 48 h. ib-lk, inclusion body-like structure. This structure was seen in close proximity with the endoplasmic reticulum (fig. a, arrows) and was positioned close to the nucleus (n) and the kinetoplast (k) in the posterior part of the cell body (fig. b) or more commonly at the anterior end, close to the kinetoplast and the basal body (fig. c). The inclusions body is not surrounded by a membrane unit (fig. d). bb, basal body; f, flagellum. Bar: 1 μm.

### 3.3. α-Tubulin hyperacetylation causes a halt in the cell cycle progression of epimastigotes

We performed growth curves of *Tc*ATAT over-expressing epimastigotes in the absence and presences of tetracycline. (Figure 5A). A growth impairment was observed after 48 h.p.i. but no differences in viability (measured with Erythrocin B staining) were observed along the entire growth curve (Supplementary Figure S5A). We have previously ruled out any undesired effect of the tetracycline treatment (Ritagliati et al., 2015a). We also quantified epimastigotes’ motility using CASA software and observed less motile parasites when epimastigotes were induced for 48 h. (Supplementary Figure S5B). Over-expression was also verified with rabbit polyclonal anti-ATAT antibodies in 24 and 48 h.p.i cells, when the *Tc*ATAT labeling was particularly strong and excluded from the nucleus and kinetoplast (Figure 5B). After 24 h.p.i. parasites that appear to have two nuclei start to accumulate (Zoom in Figure 5B, yellow arrow heads indicate parasites with an aberrant DNA content). The cell cycle progression of *Tc*ATAT overexpressing epimastigotes was analyzed by flow cytometry with Propidium iodide (PI) staining for 24 h.p.i in synchronized epimastigotes (Figure 6A). As expected, in the absence of tetracycline we observed that the cell cycle progressed normally. At time point 0 the main peak corresponds to the parasites on G1 phase of the cell cycle (∼60% of the total) that is, parasites with the DNA content corresponding to one nucleus. A second minor peak represents the parasites in G2/M phase (∼30%), which corresponds to epimastigotes with the double of DNA content, including those on cytokinesis. In the valley between the two peaks are the cells on S phase (∼10%). Then, at time point 6 hours the count of parasites in S phase starts to increase and at 12 hours more than half of the parasites are in G2/M phase. Finally, at 24 hours the parasites return to G1 phase. When tetracycline is added the peak of cells in S phase doubles at 12 h.p.i. Furthermore, G2/M peak increased about 20% at 24 h.p.i., suggesting that the *Tc*ATAT overexpression results in an arrest of cells that are dividing and cells that have duplicated their DNA (Figure 6B). This halt in the cell cycle progression was also observed by Scanning Electron Microscopy (Figure 6C). When parasites are induced with tetracycline for 24 hours, ∼30% of the cells have two flagella and it appears that there is an impairment in the cleavage furrow progression and cytokinesis (white arrows, Figure 6C g -i) that correlates with the higher proportion of cells in G2/M phase.

**Figure 5:**
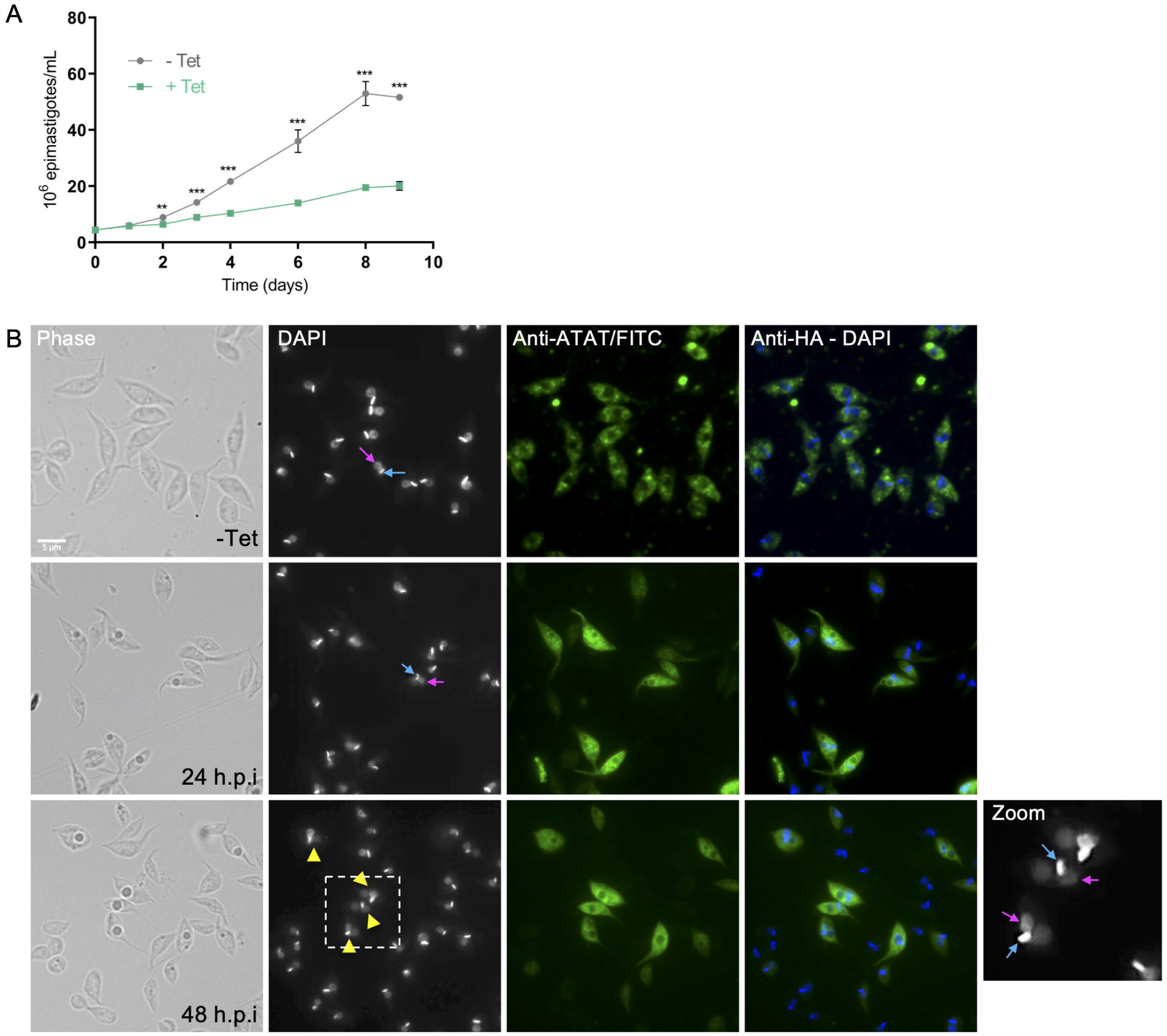
ATAT-HA over-expression negatively impacts on epimastigotes growth. **(A)** Growth curve of Dm28c p*Tc*INDEX-GW ATAT-HA epimastigotes in the absence (grey circles) and presence (green squares) of 0.5 μg/ml tetracycline for 9 days. ** p<0.005, ***p<0.001 (Student’s t-test). **(B)** Immunolocalization of *Tc*ATAT, in Dm28c p*Tc*INDEX-GW ATAT-HA epimastigotes in the absence and presence of 0.5 μg/ml tetracycline for 24 and 48 h. Bar: 5 μm. DAPI was used as nucleus and kinetoplast marker. The light blue arrow indicates the kinetoplast and the pink arrow indicates the nucleus. Yellow arrowheads indicate parasites with an aberrant DNA content.

**Figure 6:**
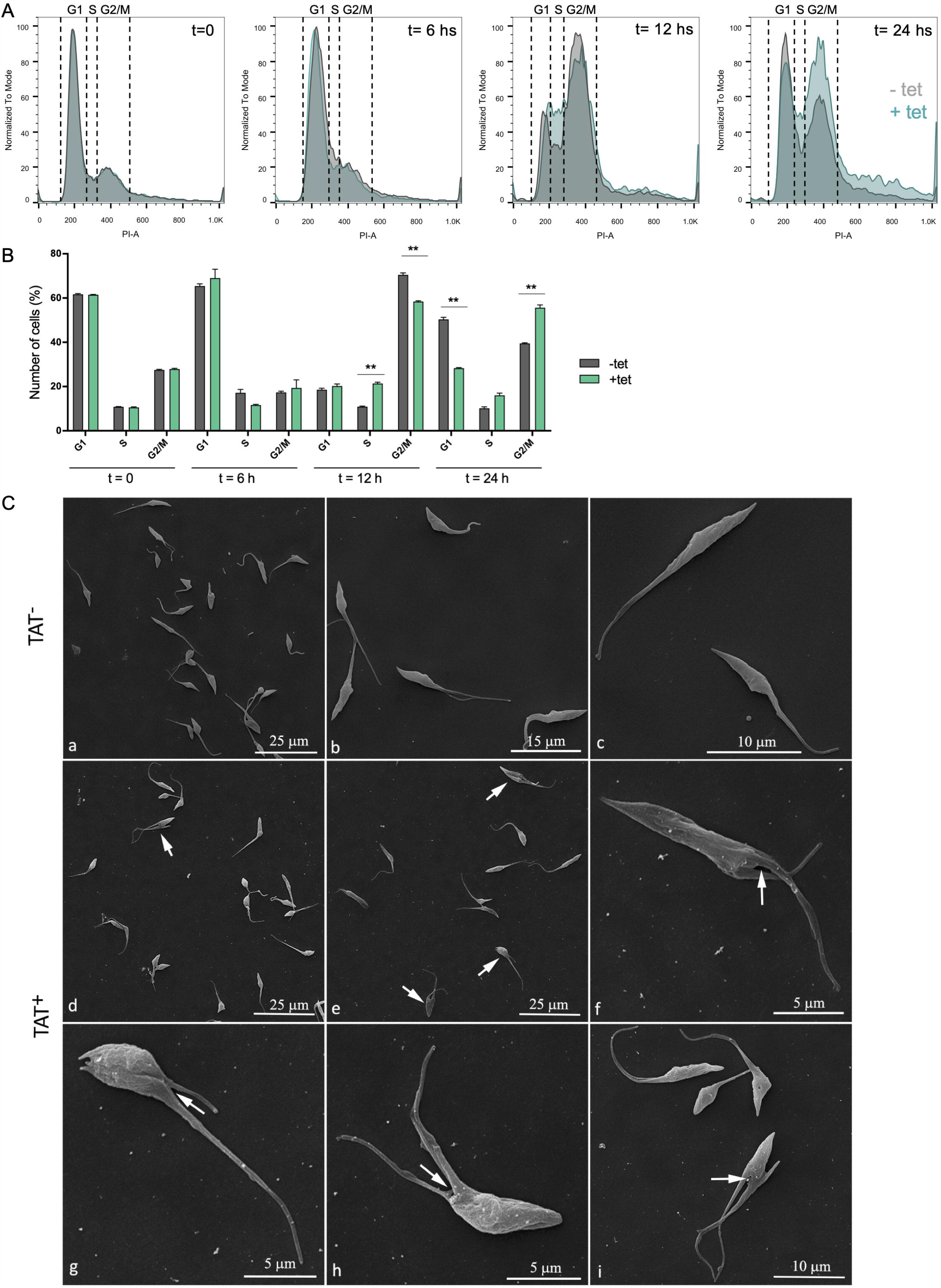
Hyperacetylation alters the cell cycle progression. **(A)** Flow cytometry analysis of synchronized Dm28c p*Tc*INDEX-GW ATAT-HA epimastigotes in the absences (grey) and presence (green) of 0.5 μg/ml tetracycline at different time points. Histograms are plotted as the normalized number of events vs. propidium iodide absorbance (PI-A). **(B)** Bar graph with the percentages of cells in the different phases of the cell cycle. **p < 0.005, ***p < 0.001 (Student’s t test). **(C)** Images obtained by Scanning Electron Microscopy (SEM) of Dm28c p*Tc*INDEX-GW ATAT-HA. Uninduced epimastigotes (TAT-) (figs. a-c). Parasites induced with 0.5 μg/ml tetracycline for 24 h (TAT+) (figs. d-i). Induced cells presented a phenotype that indicates cytokinesis arrest (figs. d-e, arrows) and the interruption in the progression of the cleavage furrow (figs. f-i, white arrows).

As a control, we quantified the population of epimastigotes with two nuclei and one kinetoplast 24 and 48 h.p.i in an asynchronous population. The ordered progression of the cell cycle, in which kinetoplast segregation precedes nuclear division, allows the identification of three normal states regarding nuclear/kinetoplast (N/K) content: 1N1K, 1N2K and 2N2K. Under normal conditions, most epimastigotes in a non-synchronous exponentially growing culture contain one nucleus and one kinetoplast (1N1K, usually ∼80–95%), corresponding to parasites in G1 or S phase of the cell cycle. A smaller proportion exhibits two kinetoplasts and one nucleus (1N2K ∼5%), these correspond to parasites in G2 phase or the beginning of mitosis. Finally, cells presenting two kinetoplasts and two nuclei (2N2K ∼3%) are those that have completed mitosis and are undergoing cytokinesis or ready to do so (Elias et al., 2007). Thus, the appearance of cells with abnormal N/K content is indicative of cell cycle impairment. 20-30% more parasites with 1K/2N are found in over-expressing conditions than in the uninduced control (Supplementary Figure S7).

### 3.4. Over-expression of *Tc*ATAT-HA alters acetylated α-tubulin distribution and causes modifications on mitochondrion ultrastructure

Parasites with over-expression of TcATAT-HA presented alterations on acetylated α-tubulin distribution observe with anti-acetylated α-tubulin antibodies. Part of the population accumulates acetylated α-tubulin around the kinetoplast (Figure 7A, yellow arrowheads) and in some parasites it is surrounding the inclusion body-like structure (Figure 7A, white arrowheads). Trypanosomatids have a single and ramified mitochondrion with the kDNA concentrated in the kinetoplast. The kinetoplast is connected to the basal body that nucleates the flagellum, that are both MT-containing structures. Since the basal body is linked to the kinetoplast by the tripartite attachment complex (TAC) (Kaser et al., 2014), we decided to investigate the mitochondrial morphology and ultrastructure in *Tc*ATAT-HA over-expressing cells by TEM and using Mitotracker Orange CMTMRos. In induced epimastigotes, cristae swelling was seen in the kinetoplast region and also in the mitochondrial branches (Figure 7B, white arrows in a and b). Moreover, sometimes cells presented a kinetoplast containing multiple networks that are very condensed (Figure 7B, white arrow in c) indicating kinetoplast division impairment. Images obtained by TEM confirmed this hypothesis since overexpressing parasites presented duplicated kDNA that did not suffer scission and was seen associated to a single basal body (Figure 7C, black arrowheads heads in b), differently to what was observed in the uninduced condition where cells contained two basal bodies (Figure 7C, black arrowheads in a). Furthermore, TEM images revealed that when the kDNA duplicated, but the kinetoplast did not divide, the network became curved, folded over itself, thus acquiring a round shape with an atypical condensation. The kinetoplast shape also changed from disk to a round format (Figure 7C, c-f). Such kDNA alterations were also observed by confocal microscopy in *Tc*ATAT-HA over-expressing parasites stained with Mitotracker and DAPI. Parasites presented a single kinetoplast with duplicated kDNA and two nuclei (Figure 7D, upper panel), as well an arched kDNA (Figure 7D, lower panel).

**Figure 7:**
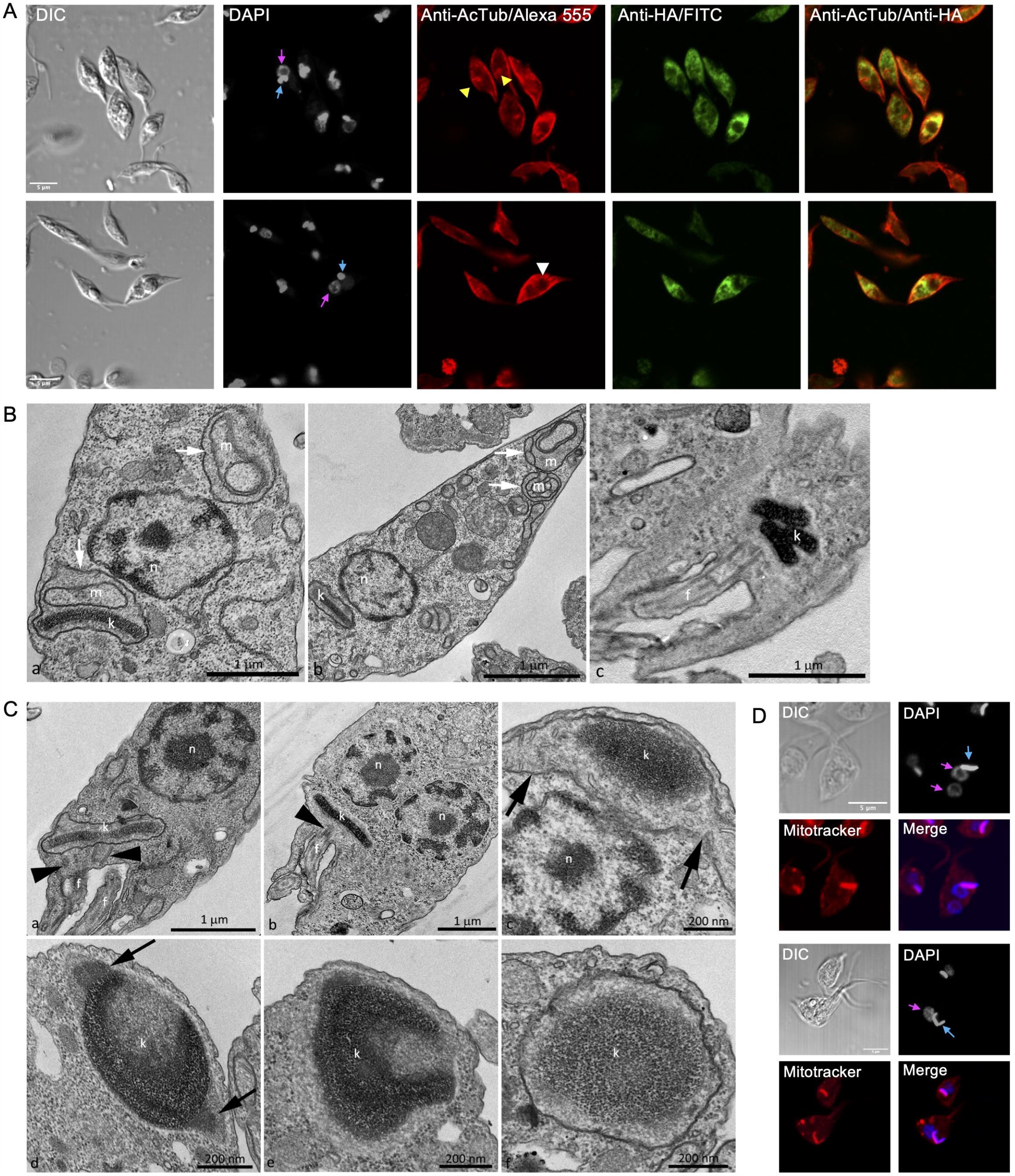
Over-expression TcATAT-HA causes phenotypic alterations in acetylated α-tubulin distribution and on mitochondrion ultrastructure of epimastigotes. **(A)** Immunolocalization of ATAT-HA with rat monoclonal anti-HA antibodies and mouse monoclonal anti-acetylated α-tubulin (anti-AcTub) of Dm28c p*Tc*INDEX-GW ATAT-HA epimastigotes induced with 0.5 μg/ml tetracycline for 24 h. Bar: 5 μm. DAPI was used as nucleus and kinetoplast marker. The light blue arrow indicates the kinetoplast and the pink arrow indicates the nucleus. Yellow arrowheads indicate accumulation of acetylated α-tubulin around the kinetoplast and the white arrowhead indicates accumulation of acetylated α-tubulin around the inclusion body-like structure. **(B)** Transmission Electron Microscopy of Dm28c p*Tc*INDEX-GW ATAT-HA epimastigotes induced with 0.5 μg/ml tetracycline for 48 h. Parasites presented alterations in the mitochondrial branches and at the kinetoplast region, especially cristae swelling (figs. a and b, white arrows). Parasites presenting a kinetoplast with multiple and electrodense networks were also observed (fig. c) f, flagellum; k, kinetoplast; m, mitochondria; n, nucleus. Bar: 1 μm. **(C)** Transmission Electron Microscopy of Dm28c p*Tc*INDEX-GW ATAT-HA. In uninduced epimastigotes the replicated kDNA is contained in a kinetoplast associated to two basal bodies (fig. a, black arrowheads). Epimastigotes induced with 0.5 μg/ml tetracycline for 48 h presented atypical characteristics (figs. b-f). In this case, the replicated kDNA is contained in a kinetoplast associated to a single basal body (fig. b, black arrowhead), which result is kinetoplast division impairment in a cell with two nuclei. The kinetoplast region is continuous with mitochondrial branches (fig. c, black arrows). The kDNA replication occurs during the S phase when the antipodal sites contain proteins involved in this process (fig. d, black arrows). Since the kDNA replicates, but the kinetoplast does not divide, the network curves and folds over itself, becoming round and presenting an atypical topology. The kinetoplast shape also changes its format from disk to round (figs. d-f). f, flagellum; k, kinetoplast; n, nucleus. Bars = 1 μm (*a* and *b*), 200 nm (*c-f*). **(D)** Dm28c p*Tc*INDEX-GW ATAT-HA epimastigotes induced with 0.5 μg/ml tetracycline for 48 h and stained with Mitotracker CMTMRos. DAPI was used as nucleus and kinetoplast marker. The light blue arrow indicates the kinetoplast and the pink arrow indicates the nucleus. The upper panel shows a parasite with a kinetoplast containing a duplicated kDNA and two nuclei, while the lower panel shows a

## 4. Discussion

In this study, we address the biological relevance of acetylated α-tubulin in the protozoan pathogen *T. cruzi*. More than 30 years ago it was described that acetylated α-tubulin was the mayor isotype present in *T. brucei* (Schneider et al., 1987; Sasse and Gull, 1988) and *T. cruzi* (Souto-Padron et al., 1993) subpellicular MTs and flagellar axoneme, but the significance of this finding has not been unraveled yet. We have identified and characterized the acetyltransferase responsible for mediating K40 α-tubulin acetylation in *T. cruzi, Tc*ATAT, and shown that its over-expression conduces to a hyperacetylation of α-tubulin that severely affects the normal progression of the cell cycle in epimastigotes. ATAT-HA over-expression also confers epimastigotes resistance to Oryzalin, a depolymerizing drug that targets α-tubulin. Dinitroaniline herbicides such as oryzalin, which was shown to depolymerize plant cell microtubules (Morejohn et al., 1987), also disrupt the microtubules of several protozoa including *Tetrahymena* (Stargell et al., 1992) and parasites such as *Leishmania spp*. (Chan et al., 1991), *Entamoeba spp*. (Makioka et al., 2000), *Cryptosporidium parvum* (Armson et al., 2002), *Toxoplasma gondii* (Stokkermans et al., 1996), *Angomonas deanei* and *Strigomonas culicis* (Catta-Preta et al., 2015). Interestingly, sensitivity of *T. cruzi* to Orlyzalin is significantly higher than for other protists where it was studied, what can be explained by the presence of a L at position 267, instead of a V or I found in *Toxoplasma gondii* (Shaw et al., 2000) and *Tetrahymena thermophila* (Dostál and Libusová, 2014) respectively, among other point mutations found in *T. cruzi* α-tubulin.

Acetylated α-tubulin has been associated with stable structures in eukaryotic cells, localizing to primary cilia, midbodies, centrioles and subsets of cytoplasmic microtubules in 3T3 and HeLa cells (Piperno et al., 1987) and to flagella axonemes, basal bodies and cytoplasmic microtubules radiating from the basal bodies in *Chlamydomonas reinhardtii* (L’Hernault and Rosenbaum, 1985). In *T. brucei* and *T. cruzi* acetylated α-tubulin is distributed widely throughout all microtubular arrays (Sasse and Gull, 1988; Souto-Padron et al., 1993). This post-translational modification appears to occur during or immediately after microtubule polymerization, and the deacetylation process correlates with depolymerization (Sasse and Gull, 1988). The fact that we observed a resistance to an α-tubulin depolymerizing drug when MTs are hyperacetylated suggests that there is a clear link between acetylation and stabilization in *T. cruzi* as reported in other organisms. Subpellicular microtubules that compose the trypanosomatid cytoskeleton are quite stable structures, but our results suggest that a fine regulation in tubulin polymerization/depolymerization is necessary for the correct progression of the cell cycle and protozoan division.

Early electron microscopy revealed distinct subcellular sites from which microtubules appeared to emanate which were named ‘microtubule-organizing centers’ in eukaryotes (MTOCs). Since then, the exact nature of MTOCs has remained unclear. Microtubules have an inherent structural polarity, with a dynamic plus end and a comparatively stable and slow growing minus end. These characteristics of microtubule minus ends can be influenced *in vivo* by an association with a MTOC, that can be broadly defined as sites for microtubule nucleation, stabilization, and/or anchoring (Sanchez and Feldman, 2017). Not much is known about MTOCs in trypanosomatids apart from the fact that subpellicular microtubules have uniform spacing over the entire parasite, presenting their minus ends oriented toward the anterior pole of the cell, the region where the single-copy organelles division starts (Wheeler et al., 2019). Trypanosomes have γ-tubulin and γ-tubulin ring complex proteins, but unfortunately their localization or interrogation of function has not led to the definition of the sites of individual microtubule nucleation within the subpellicular array (Zhou and Li, 2015). We observed that ATAT is concentrated in perinuclear spots in epimastigotes and amastigotes, and that it accumulates mainly in the anterior region when over-expressed. These results as well as the fact that α-tubulin acetylation occurs immediately after microtubules polymerization may suggest that ATAT stabilizes the microtubules that form the subpellicular corset when they are nucleating in the MTOC.

The *T. cruzi* cell cycle is characterized by a coordinated duplication of nuclear and kinetoplast DNA during the S phase. After kDNA replication, the kinetoplast assumes a more elongated disk shape and segregates at the beginning of the G2 phase. At this point cells present two basal bodies, both linked to the kDNA network (Elias et al., 2007). TEM analyses revealed that many cells showed an elongated kinetoplast, indicating that the kDNA replication occurred, but not the network scission. This is related to hyperacetylation that also caused a marked halt in G2/M phase of the cell cycle as determined by flow cytometry. A phenotype related to *Tc*ATAT over-expression is the impaired ingression of the cleavage furrow, resulting in a defect in cytokinesis, an observation that can be associated to an increase in the amount of acetylated α-tubulin (or perhaps to a decrease in the amount of non-acetylated α-tubulin available). The blocked cytokinesis resembles the phenotype of *T. brucei* GTPase Arl2 mutants. Arl2 orthologues in mammals are mitochondrial proteins but in *T. brucei* it appears to be a cytoskeletal protein. Knockdown and over-expression of *Tb*Arl2 modulate the levels of acetylated α-tubulin and inhibits cytokinesis and cleavage furrow progression similar to ATAT-HA over-expression (Price et al., 2010). Furthermore, in *T. brucei* cytokinesis proceeds from the anterior end to the posterior end, with the cleavage furrow starting at the distal tip of the new Flagellum Attachment Zone (FAZ) and proceeding along a fold in the cell. The furrow placement relative to the old and new FAZ guarantees correct inheritance of the basal body, kinetoplasts, and flagellar pocket complexes, but the nuclei must be positioned correctly. Finally, the furrow resolves to a single point of connection between the posterior of one daughter cell and the side of the other daughter. This narrow cytoplasmic bridge can persist while the daughter cells restart the cell cycle, although it is normally resolved (Wheeler et al., 2019). In conclusion, furrow ingression must require some rearrangement of the microtubule array. It is quite possible that an increase in α-tubulin acetylation, as a consequence of ATAT-HA over-expression, somehow promotes the stabilization between kDNA and the basal body (which are also composed by MTs) thus impairing MTs rearrangements and cytokinesis. Basal body replication can be impaired in cells over-expressing ATAT-HA, since protozoa containing duplicated kDNA network, a single basal body and only one flagellum were observed. Unfortunately, we still do not know much about the cell cycle checkpoints of *T. cruzi* epimastigotes, what would enable a deeper discussion about the observed phenomenon.

A refringent button-like structure was visible by optic microscopy that started to grow in a time-dependent manner after induction of ATAT-HA over-expression with tetracycline. This round structure contains ATAT-HA, forming an insoluble and tridimensional structure that remains associated with isolated cytoskeletal and flagellar fractions. This atypical structure is electrodense and is observed by TEM most of the time in the anterior region, close to the nucleus and kinetoplast. It is not delimited by a membrane unit, which suggests that it could be a cumulus of protein, rich in *Tc*ATAT-HA, reminiscent to an inclusion body. Inclusion bodies are aggregates of misfolded protein known to occur in eucaryotic cells, for example during neurodegenerative disorders (Chung et al., 2018). Inclusion bodies are also found in bacteria as particles of aggregated protein (Singh and Panda, 2005). To our knowledge there are no reports of inclusion bodies in trypanosomatids occurring as a consequence of over-expression of exogenous proteins. It is also worth mentioning that we have use this over-expression systems for different proteins and never observed this phenotype (Ritagliati et al., 2015b, 2015a; Alonso et al., 2016; Tavernelli et al., 2019). We believe that these inclusion-body like structures are not occurring due to protein misfolding but as a consequence of the accumulation of ATAT-HA in a specific region the cytoskeleton. Suggestively, we also observe an accumulation of acetylated α-tubulin around the kinetoplast and the inclusion body-like structure at the anterior region, where in normal conditions *Tc*ATAT is proposed to acetylate the microtubes as they polymerize. It is proposed that *T. cruzi* kinetoplast division is similar to that described to *Crithidia fasciculata* t (Ferguson et al., 1994; Liu et al., 2005) and since the kinetoplast is part of the single mitochondrion, it is suggested that trypanosomatid’s mitochondrion could start to segregate in the kinetoplast region (Ramos et al., 2011), which correlates with our observations. We propose that hyperacetylation impairs the division of the kDNA, given that the kinetoplast divides in coordination with the basal body. Cytoskeletal elements such as the flagellum, FAZ, flagellar pocket and the subpellicular microtubule array, all need to be duplicated and segregated in a coordinated manner in relation to the nuclear and kinetoplast cycles. The basal body in trypanosomes is the master organizer for the surrounding cytoskeleton, membranous structures, and organelles. Regulation of the basal body maturation, biogenesis, segregation and positioning is vital to ensure the shape and form of subsequent daughter cells (Elias et al., 2007). Further studies are required to determine the exact role of α-tubulin acetylation in the division of the kinetoplast and the basal body.

Oliveira Santos et al, described the effect of Trichostatin A (TSA), a deacetylase inhibitor in *T. cruzi*. They report that one of the main effects of TSA treatment is α-tubulin hyperacetylation, which induced microtubule cytoskeleton reorganization. They observed the presence of parasites with replicated kDNA, associated with basal bodies, but an incomplete cytokinesis. They also reported a higher number of protozoa in G2/M phase of the cell cycle and polynucleated cells with an aberrant phenotype (Santos et al., 2018). These results are similar to those observed in ATAT-HA over-expressing cells. Probably TSA treatment is targeting several deacetylases in *T. cruzi*, so it could promote the hyperacetylation of other proteins besides α-tubulin, but the cytoskeletal remodeling seems to be linked to α-tubulin acetylation.

Our study is the first report of the ATAT/MEC-17 homologue in trypanosomatids. Besides ATAT/MEC-17 itself, no other substrates than α-tubulin have been reported for this family of lysine acetyltransferases to date. Mammalian ATAT has been shown to localize to the lumen of MTs, where it exerts its KAT activity *in vitro* (Szyk et al., 2014). *Tc*ATAT-HA tight association with MTs could also be due to the same luminal localization but this needs further corroboration. Our results suggest that a precise amount of acetylated/non-acetylated α-tubulin is necessary for the correct kinetoplast division and assembly/disassembly of the basal body and the flagellum in epimastigotes. Further experiments are needed to determine the possible effect of α-tubulin hyperacetylation over the other stages of *T. cruzi* life cycle.

## 6. Author Contributions

Conceived and designed the experiments: VLA, MC, MCM, CSG, ES. Performed the experiments: VLA, MC, CSG, GMP, MEC, AP, LET. Wrote the manuscript: VLA, MCM and ES. All authors contributed with data analysis, share the responsibility related to the accuracy of the work, revised the manuscript, and approved its final version.

## 7. Funding

This work was supported by Agencia Nacional de Promoción Científica y Tecnológica, Ministerio de Ciencia, Tecnología e Innovación Productiva, Argentina [PICT 2017–1978], Universidad Nacional de Rosario [PIP 1BIO490] and Research Council United Kingdom [MR/P027989/1] and also by Conselho Nacional de Desenvolvimento Científico e Tecnológico (CNPq) and Fundação de Amparo à Pesquisa do Estado do Rio de Janeiro (FAPERJ).

### 8. Acknowledgments

We would like to thank Rodrigo Vena, Dolores Campos, Romina Manarin and Mara Ojeda for their technical assistance in confocal microscopy, cells and parasites culture and flow cytometry respectively. Also, we are grateful to Ariel Silver for the anti-PAR2 antibodies and Carla Ritagliati for the assistance with motility measurements using the CASA Hamilton equipment.

