## Supplementary Figure for "Alpha-tubulin acetylation in *Trypanosoma cruzi*: a dynamic instability of microtubules is required for replication and cell cycle progression"

Supplementary Material

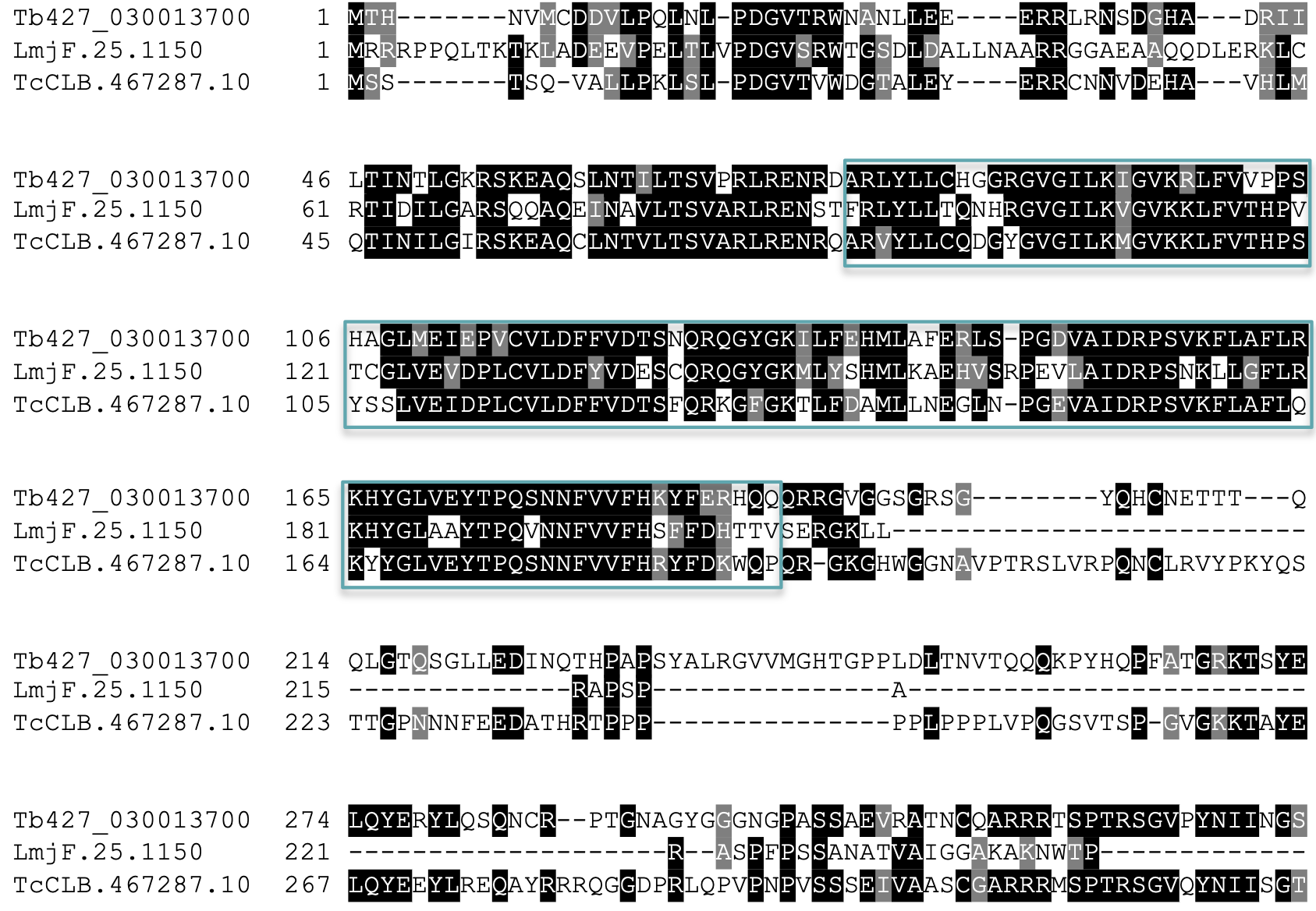

**Supplementary Figure 1.** Multiple sequence alignment of *Tc*ATAT (TcCLB.467287.10) and its homologues in *Trypanosoma brucei* (Tb427_030013700) and *Leishmania mayor* (LmjF.25.1150) using T-coffee server and colored with Boxshade. The acetyltransferase domain is boxed in green.

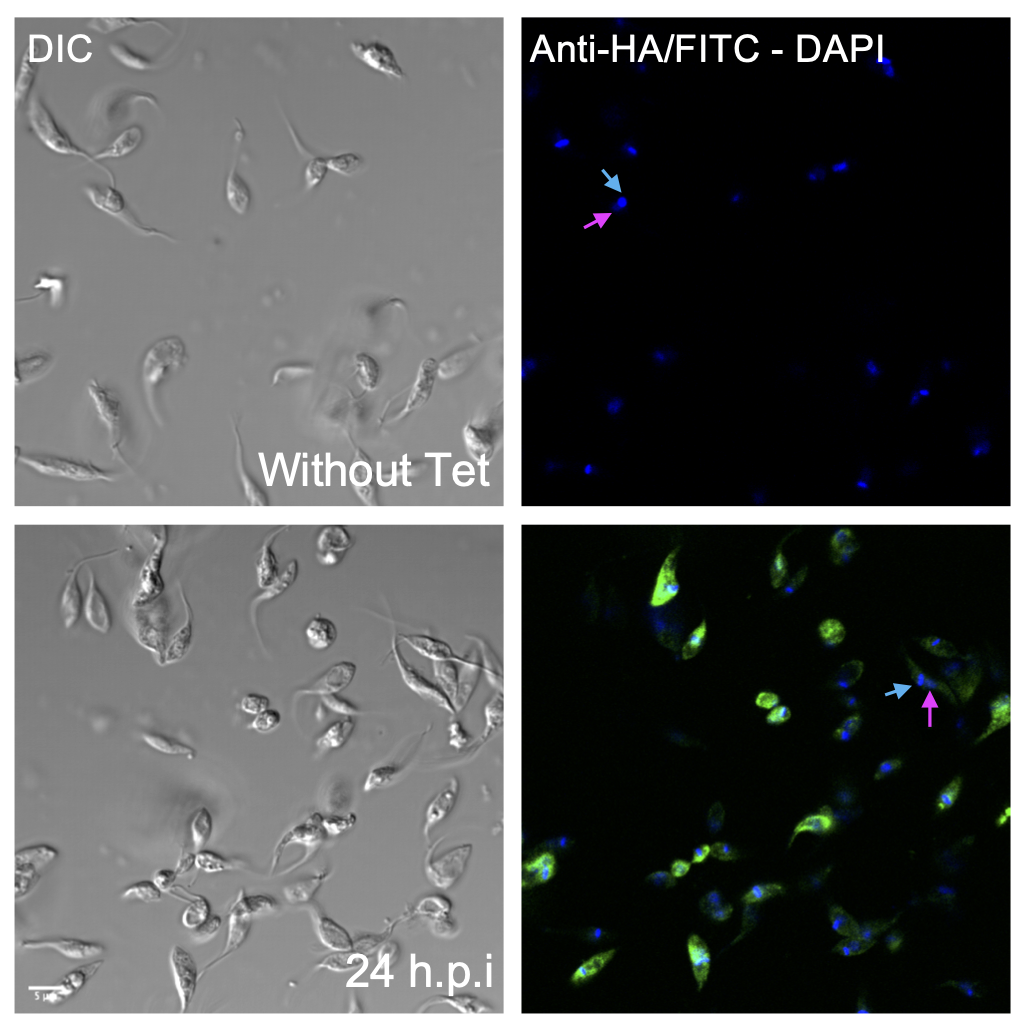

**Supplementary Figure 2.** Immunolocalization of ATAT-HA with rat monoclonal anti-HA antibodies in Dm28c p*Tc*INDEX-GW ATAT-HA epimastigotes uninduced (Without tet) and induced with 0.5 μg/ml tetracycline 24 hours post-induction (h.p.i). Bar: 5 μm. DAPI was used as nucleus and kinetoplast marker. The light blue arrow indicates the kinetoplast and the pink arrow indicates the nucleus.

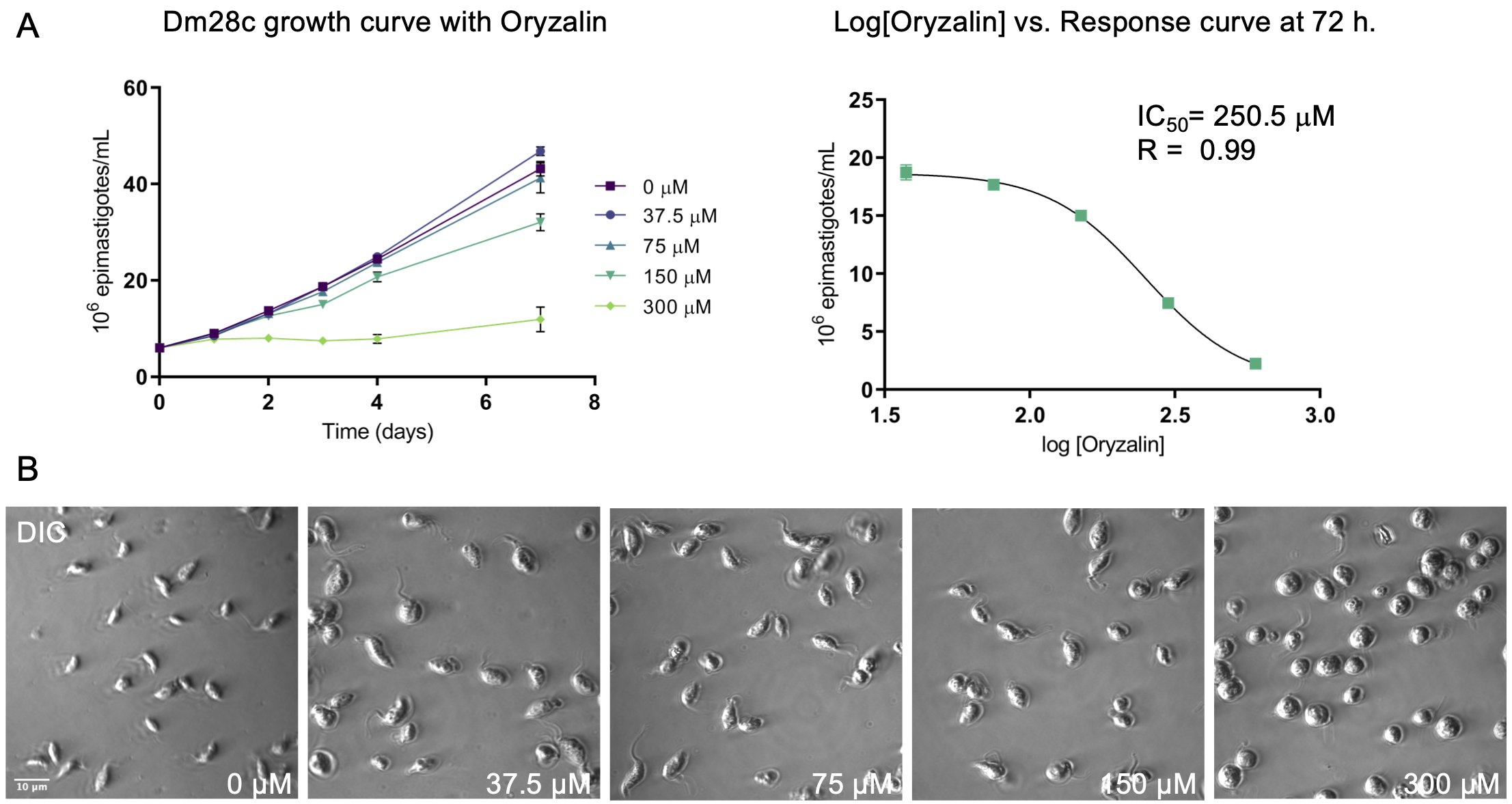

**Supplementary Figure 3. (A)** Growth curve of Dm28c epimastigotes in the presence of increasing concentrations of Oryzalin (0-300 μM) (left panel) and number of parasites *versus* the log [Oryzalin] at 72 h (right panel). The latter plot was fitted with the non-parametric regression log(inhibitor) vs. response -Variable slope (four parameters) in GraphPad Prism version 8.0 to obtain the IC_50_ value and the R^2^ of the fit. **(B)** DIC images of the morphological changes in Dm28c epimastigotes with different concentrations of Oryzalin at 72 hours. Bar = 10 μm.

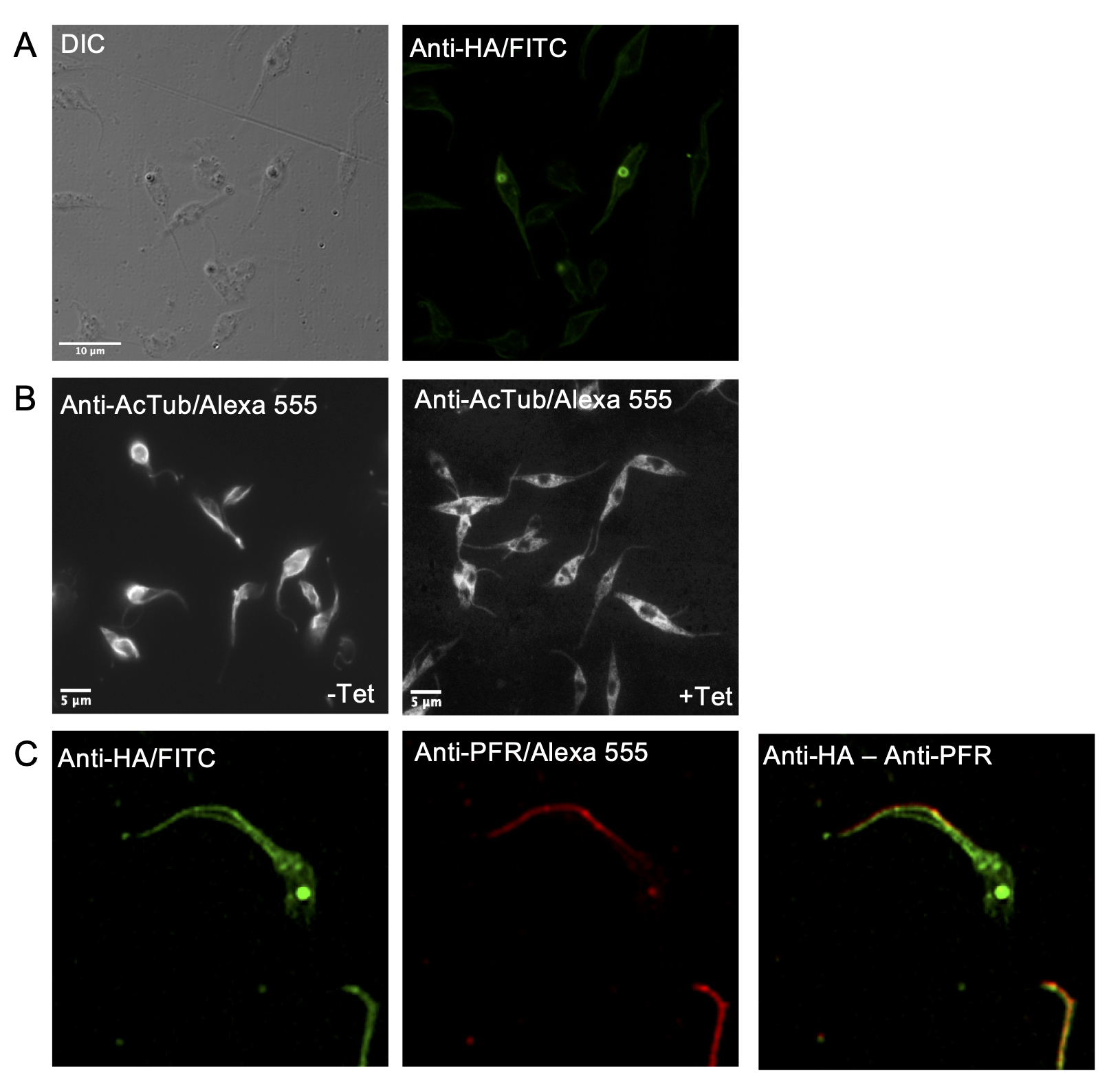

**Supplementary Figure 4. (A)** Immunolocalization of ATAT-HA with rat monoclonal anti-HA antibodies **in** isolated cytoskeletons of Dm28c p*Tc*INDEX-GW ATAT-HA epimastigotes induced with 0.5 μg/ml tetracycline for 24 h **(B)** Immunolocalization of acetylated α-tubulin with mouse monoclonal anti-acetylated α-tubulin (anti-AcTub) in isolated cytoskeletons of Dm28c p*Tc*INDEX-GW ATAT-HA epimastigotes uninduced (-Tet) and induced with 0.5 μg/ml tetracycline for 24 h (+ Tet). **(B)** Immunolocalization of ATAT-HA with rat monoclonal anti-HA antibodies and mouse polyclonal anti-paraflagellar rod 2 from *T. cruzi* (anti-PFR) in isolated cytoskeletons of Dm28c p*Tc*INDEX-GW ATAT-HA epimastigotes induced with 0.5 μg/ml tetracycline for 24 h.

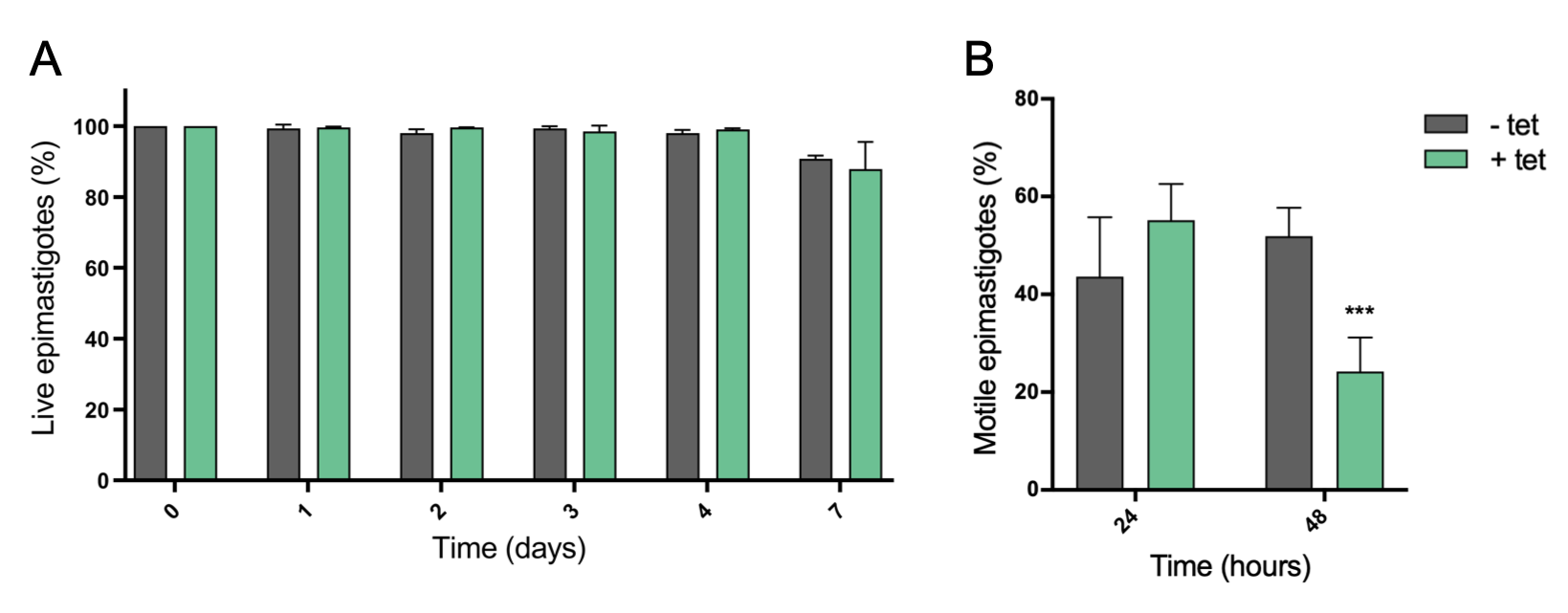

**Supplementary Figure 5. (A)** Viability (% of live epimastigotes) of Dm28c p*Tc*INDEX-GW ATAT-HA epimastigotes uninduced (grey bar, -tet) and induced with 0.5 μg/ml tetracycline (green bar, +tet) determined by counting live cells with a hematocytometer using Erythrosin B staining for 7 days. **(B)** Epimastigotes movements were examined using the computer-assisted semen analysis (CASA) system (Microptic, SCA evolution). The mean path velocity (VAP, μm/sec) was plotted at 24 and 48 h.p.i. without (grey bar, -tet) and with 0.5 μg/ml tetracycline (green bar, +tet). Experiments were performed in triplicates.

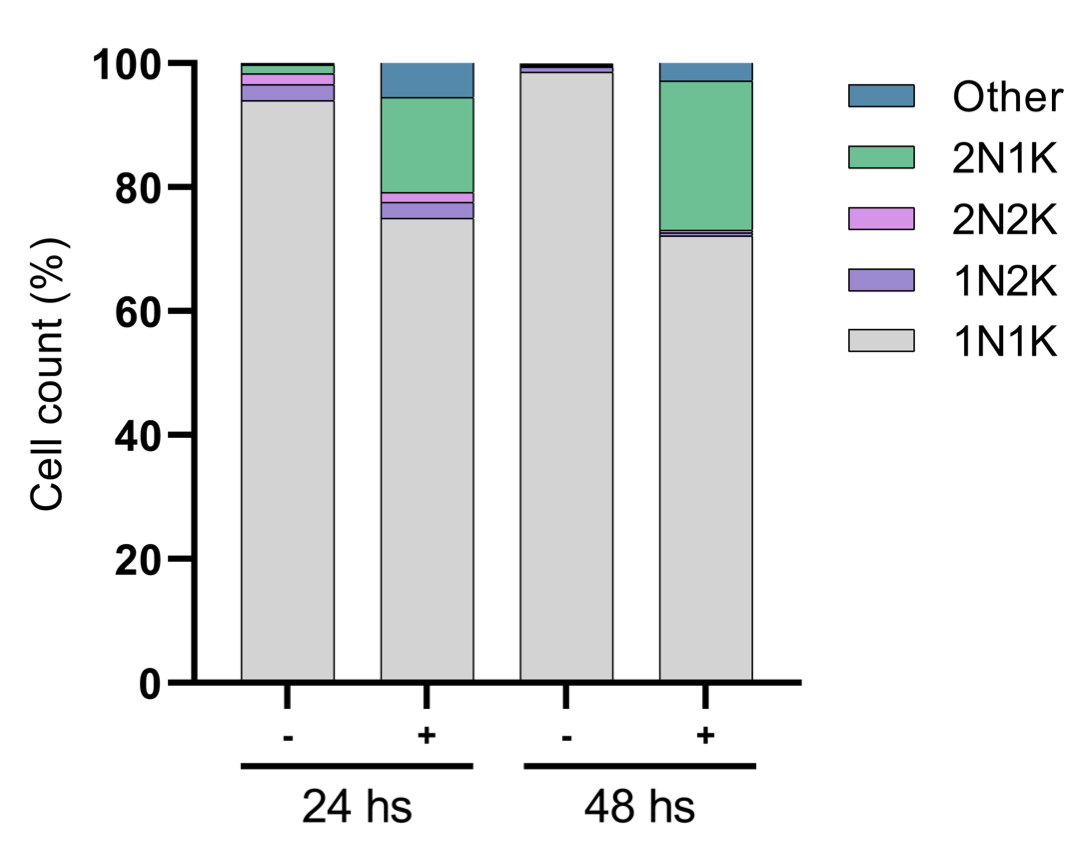

**Supplementary Figure 6.** Nucleus/Kinetoplast content (N/K) of Dm28c p*Tc*INDEX-GW ATAT-HA epimastigotes cultures in the absence (-) or presence (+) of 0.5 μg/ml tetracycline at different time points (24 and 48 hours). Data from three independent experiments were considered in the analysis (n = 300 cells for each column).
